# FAMSA2 enables accurate multiple sequence alignment at protein-universe scale

**DOI:** 10.1101/2025.07.15.664876

**Authors:** Adam Gudyś, Andrzej Zielezinski, Cedric Notredame, Sebastian Deorowicz

## Abstract

We introduce FAMSA2, an algorithm that produces high-accuracy multiple protein sequence alignments with unprecedented speed. Across structural, phylogenetic, and functional benchmarks, FAMSA2 matches or exceeds the accuracy of state-of-the-art tools while running up to 400 times faster and outperforming the fastest existing aligners. The ability of FAMSA2 to process 12 million sequences in 40 minutes on a 64 GB RAM workstation enables evolutionary and structural analyses at previously unattainable scales.

## Main text

The output of sequencing has increased a billion-fold in less than 30 years^1^, fueling initiatives such as the Earth BioGenome Project^2^ and MGnify^3^, which aim to catalog all genomic information on the planet. As a result, protein sequence databases will soon contain billions of entries. To keep pace, sequence similarity search tools able to process this amount of data have been developed (DIAMOND^4^, MMseqs2^5^), enabling the discovery of hundreds of thousands of novel protein families^6^. In parallel, the AlphaFold revolution^7^ and the release of over 200 million protein structures^8^ have driven the development of structure-based clustering tools (i.e., FoldSeek^9^), further enriching protein family diversity. Multiple sequence alignment (MSA) tools—crucial for inferring evolutionary relationships, protein structure, and phylogeny—lag, however, behind the rapidly growing protein data. The most accurate aligners for large datasets, Muscle5^10^ and T-Coffee’s regressive method^11^, scale poorly beyond thousands of sequences, with accuracy degrading as dataset size and noise increase^12^. As protein families grow towards millions of members, new alignment approaches free of these limitations are urgently needed.

We address these issues with FAMSA2, a progressive MSA algorithm (**Fig. 1a**) that aligns millions of sequences in minutes, efficiently utilizing modern hardware. FAMSA2 reaches or exceeds the accuracy of state-of-the-art tools, exhibits greater throughput than the fastest existing methods, shows superior robustness to non-homologous contamination, and provides usability features not offered by other approaches, nor its predecessor^13^ (**Fig. 1b**). FAMSA2 introduces a stochastic medoid tree algorithm that removes practical barriers to large-scale alignment by generating stable guide trees in *O*(*M N* log*N*) time—where *N* is the number of sequences and *M* the maximum subtree size—with accuracy comparable to default *O*(*N*^2^)-time single-linkage trees (**Fig. 1c**). This guide tree approximation is supported by algorithmic advances that improve both alignment accuracy and throughput, including: (i) a longest common subsequence (LCS)-based dissimilarity measure resilient to sequence divergence; (ii) length-based prefiltering of pairwise comparisons; (iii) multilevel parallelism at LCS calculation (**Fig. 1d**) and progressive alignment stage; (iv) parallel construction of single-linkage guide trees in linear space using Prim’s algorithm (**Fig. 1e**); and (v) dynamic refinement of partial alignments.

**Fig. 1.**
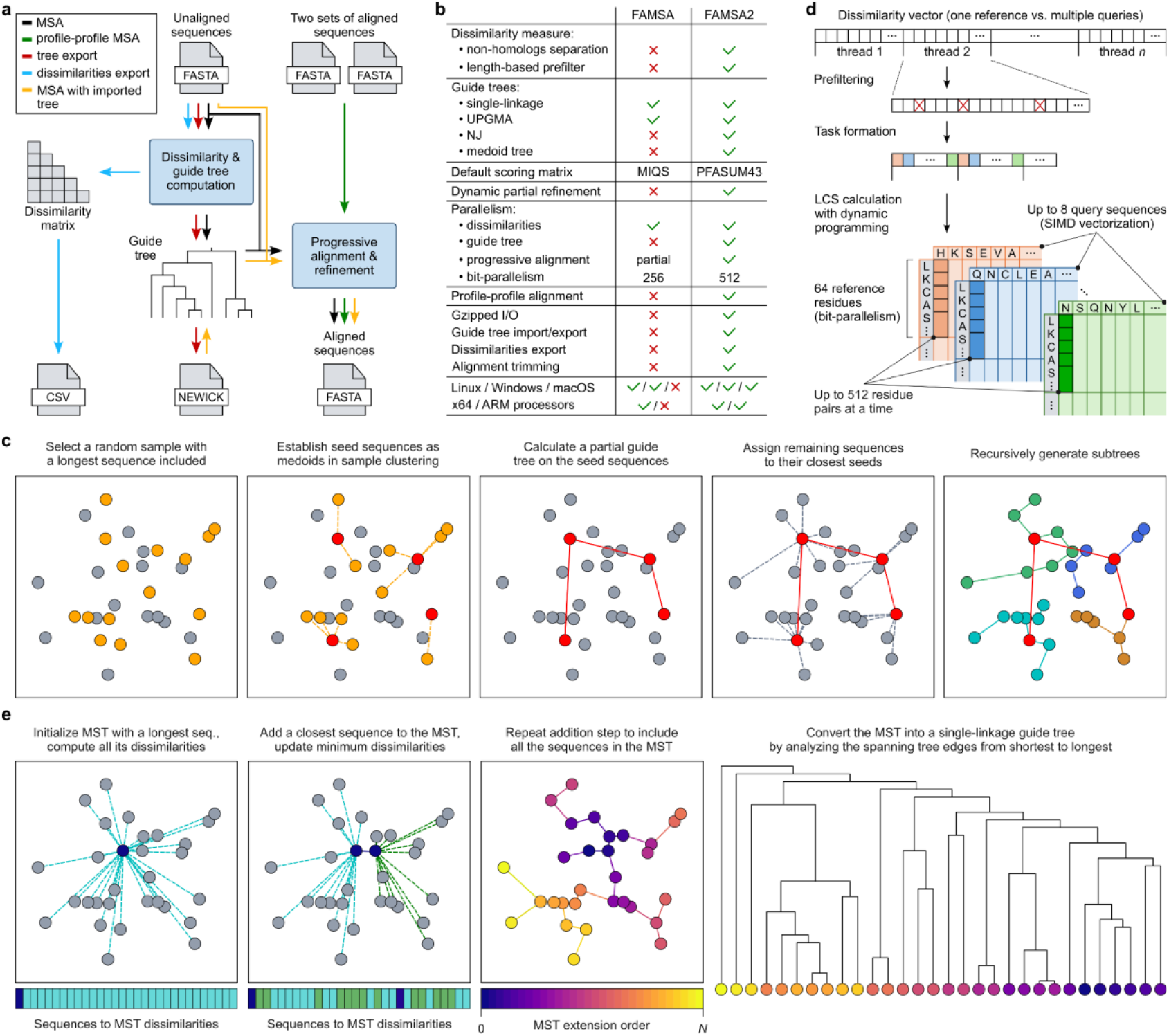
FAMSA2 algorithms and features. **a,** Usage scenarios presented as data streams. **b**, Major differences between FAMSA and FAMSA2. **c**, Medoid tree algorithm for fast guide tree approximation with reduced stochasticity in seed selection and improved stability over fully random algorithms like PartTree. **d**, Parallel dissimilarity calculation between the currently analyzed sequence (reference) and all other sequences in a set (queries) during guide tree construction. **e**, Single-linkage guide tree construction with linear-space Prim’s minimum spanning tree (MST) algorithm.

We evaluated FAMSA2 across benchmarks, assessing structural, phylogenetic, and functional alignment accuracy. The structural accuracy was measured using an updated extHomFam^13^, the largest structure-based benchmark, containing 390 protein families with up to 3 million sequences. Each family includes a subset of sequences with curated, structurally-based reference alignments, used to measure the percentage of correctly aligned residue pairs (sum-of-pairs; SP) and columns (total-column; TC) (Methods). Muscle5, T-Coffee, and Clustal Omega required 8–37 days to process this benchmark and failed at the largest ABC transporters family (**Fig. 2a, Table S1**). In contrast, FAMSA2 produced the most accurate alignments in 15 hours (SP=79.6, TC=61.8), while FAMSA2 with medoid trees required only 47 minutes to deliver results comparable to Muscle5 (>400x slower) and only slightly inferior to T-Coffee (>1,130x slower). Moreover, this fast mode of FAMSA2 surpassed Clustal Omega, MAFFT, and Kalign by more than 15 points in SP and TC (**Fig. 2a**), showing that medoid tree properties (**Extended Data Fig. 1**) translate to superior accuracy. The advantages of FAMSA2 in accuracy and throughput became more pronounced as protein family size increased (**Fig. 2b, Table S1**).

**Fig. 2.**
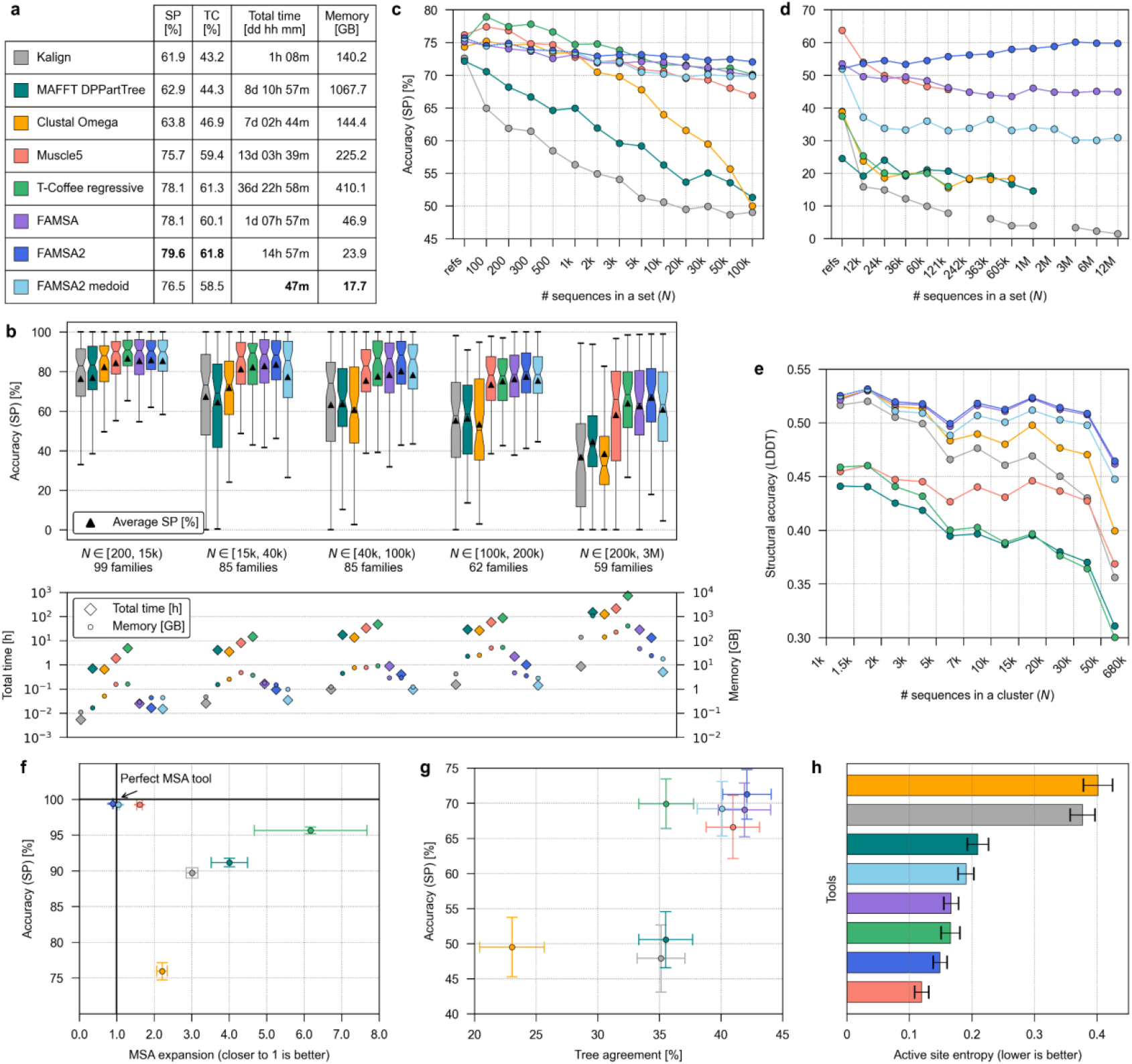
Benchmarking FAMSA2 across diverse datasets. **a**, Average accuracy, total runtime, and peak memory usage on 390 extHomFam protein families. Accuracy is shown as sum-of-pairs (SP) and total-column (TC) scores. **b**, Accuracy, runtime, and peak memory across five extHomFam family size groups. Box plots show median (centre line) with 95% confidence interval (CI, notches), mean (triangle), upper and lower quartiles (boxes), and 1.5x of range between upper and lower quartiles (whiskers). **c**, Average SP scores for 121 largest extHomFam families subsampled to contain 100–100,000 homologs. **d**, Average SP scores for mixed homologous/non-homologous datasets. **e**, Structural MSA accuracy as bootstrapped LDDT score on clusters of increasing sizes sampled from AlphaFold Database. **f**, Average phylogenetic accuracy (MSA expansion; the number of columns relative to the reference MSA) and structural accuracy (SP) on 1,860 simulated MSAs; error bars indicate the 95% CI. **g**, Average phylogenetic accuracy (tree agreement) and structural accuracy (SP) on 113 extHomFam families sampled to 100,000 sequences; error bars: 95% CI. **h**, Accuracy on 772 enzyme families with known active sites measured as an average entropy of activesite residue distribution across alignment columns; error bars: 95% CI. A summary of these benchmark results (a-h) is provided in **Extended Data Table 1**.

To further assess accuracy scaling, we analyzed the largest extHomFam families subsampled to contain from 100 to 100,000 homologs (**Fig. 2c**, Methods). FAMSA2 ranked first in accuracy for families with ≥5,000 sequences, outrunning Muscle5 from *N* ≥ 700 and T-Coffee from *N* ≥ 5,000. It also showed the most stable performance across all family sizes, with an SP score decrease of only 3.6 points, compared to 9.2 and 6.7 point-declines for Muscle5 and T-Coffee, respectively (**Fig. 2c, Table S2**). FAMSA2’s medoid trees had comparable stability and accuracy to Muscle5, while the other tools—MAFFT, Clustal Omega, and Kalign—showed lower accuracy, with SP score declines exceeding 20 points.

To investigate alignment robustness to contamination by unrelated sequences, we generated datasets containing an increasing number of mixed homologous and non-homologous sequences, scaling up to over 12 million sequences (**Fig. 2d**, Methods). Only FAMSA and FAMSA2 completed all datasets, as the others failed due to memory or time constraints (**Fig. 2d, Table S3**). FAMSA2 achieved the highest SP/TC scores across 12 out of 14 and, unlike other tools, improved its accuracy as dataset size increased. This resulted from the modified dissimilarity measure, which effectively clusters related sequences (**Extended Data Fig. 2**). FAMSA2’s medoid tree ranked third in SP/TC stability and overall accuracy, following FAMSA and FAMSA2 (**Fig. 2d**).

To address the potential limitations of extHomFam benchmark, i.e., its reliance on a small number of reference structures, we aligned ∼1,100 protein families (1,000–∼680,000 sequences) from AlphaFold Database Clusters^8^ and evaluated the resulting MSAs using the LDDT score^16^—a reference-free accuracy measure based on the local structures superposition (0–1; Methods). Both FAMSA2 variants achieved the highest average LDDT scores (0.50–0.51) (**Extended Data Table 1, Table S4**), and also had the smallest decline in accuracy when increasing dataset sizes (**Fig. 2e**). Notably, Clustal Omega (0.48) and Kalign (0.46), which ranked lower in the previous benchmarks, surpassed Muscle5 (0.43) and T-Coffee (0.40) in this structural assessment. The medoid version of FAMSA2 required 227-fold less time than Clustal Omega (**Table S5)**, the next most accurate method, while Kalign, the only algorithm with comparable execution times, delivered alignments of significantly lower quality. Overall, FAMSA2 processed the entire database of ∼25,000 families in half a day compared to an estimated ∼59 days for Clustal Omega.

Given that structural accuracy is only a proxy of evolutionary correctness, we investigated the capacity of alignments to retain phylogenetic information using structure-free experimental protocols. First, we simulated 1,860 protein families along known trees (Methods), yielding reference MSAs grounded in evolutionary history. Both FAMSA2 variants achieved the highest SP/TC scores (**Fig. 2f, Table S6**) with near-ideal expansion—the ratio of the number of columns in the tool’s MSA to those in the reference MSA (**Fig. 2f**). Next, we assessed phylogenetic signal on real protein families. In the absence of reference phylogenies, we compared maximum likelihood (ML) trees, often considered a ground truth in phylogenetic comparison, and distance-based (minimum evolution, ME) trees reconstructed from the extHomFam protein families, using topological agreement as a proxy for phylogenetic accuracy (Methods, **Table S7**). FAMSA2 achieved the highest tree agreement score of 42.1%, followed closely by FAMSA (41.9%) and Muscle5 (41.0%) (**Fig. 2g**). The FAMSA2 medoid tree ranked next with 40.1% and outperformed the other tools by 4–17 percentage points. Although the high ME-ML trees agreement stops short of demonstrating improved phylogenetic accuracy, it supports the notion of noise reduction when using FAMSA alignments in phylogeny.

Our final experiment focused on functional alignment, as structure-and phylogeny-based benchmarks may be confounded by convergent evolution when used for functional modeling. We assessed the preservation of 1,376 active sites across 772 Pfam enzyme families by measuring the entropy of the distribution of active sites’ residues across alignment columns (Methods; lower entropy indicates better preservation; **Table S8**). FAMSA2 achieved an entropy of 0.15 (second after Muscle5 (0.12)) (**Fig. 2g**) retaining on average 97% of active sites residues in the most populated columns (**Extended Data Fig. 3**). The medoid-based variant of FAMSA2 also produced accurate results (0.19) slightly surpassing MAFFT (0.21), and outperforming Kalign (0.38) and Clustal Omega (0.40), which both spread active sites across more columns (<95% of residues in the first column).

In conclusion, FAMSA2 combines high accuracy with unmatched scalability, enabling efficient exploitation of current and future multi-core architectures (**Extended Data Fig. 4**). It excels in two key scenarios: aligning massive protein families—such as the ∼3-million-sequence ABC transporter family in 5 minutes using 18 GB of RAM—and processing large collections, including ∼22,000 Pfam-A families (∼62 million sequences) in 8 hours, and 100,000 recently discovered protein families (∼20 million sequences)^6^ in ∼5 hours. The medoid tree algorithm provides further evidence on the suitability of non-biological trees for MSA reconstruction^14,15^, bringing a renewed long-term sustainability to progressive alignment. With FAMSA2, MSAs are now able to scale with the new-generation databases and family discovery tools and ready to face the challenge of entire eco-system sequencing.

## Acknowledgments

This work was supported by the National Science Centre, Poland, project DEC-2022/45/B/ST6/03032 to SD and AG. The computations were partially performed at the Poznan Supercomputing and Networking Center (grant number pl0074-02).

## Author contributions statement

SD and AG designed the study and developed FAMSA2. AG, SD, and AZ performed benchmarking. AG, AZ, CN, and SD analyzed the results and wrote the manuscript. All authors reviewed and approved the manuscript.

## Competing interests statement

The authors declare no competing interests.

## Online Methods

### FAMSA2 overview

FAMSA2 retains the two-stage progressive alignment approach of its predecessor. In the first stage, the guide tree is constructed using either our new medoid tree algorithm or a standard single-linkage clustering (other tree algorithms like UPGMA or NJ are available as options). In the second stage, the sequences are aligned into profiles (intermediate alignments) following the incorporation order indicated by the guide tree.

Beyond full MSA computation, FAMSA2 supports the following workflows: (*i*) guide tree generation, (*ii*) alignment using an external guide tree, (*iii*) dissimilarity or identity matrix calculation, and (*iv*) alignment of two input profiles (**Fig. 1a**).

#### Medoid tree

Medoid tree is a stochastic algorithm for guide tree approximation characterized by improved scalability compared to Prim’s algorithm used for the default single linkage trees. In particular, medoid trees require *O*(*M N* log*N*) time (where *M* is the algorithm parameter representing maximum subtree size), which is a significant advance over Prim’s *O*(*N*^2^) for large sequence sets.

The medoid tree algorithm is inspired by MAFFT PartTree^17,18^ and addresses its main issue, namely, the substantial randomness. PartTree performs a divisive sequence clustering through the following steps:

1. Assign all *N* sequences to the initial cluster
2. Select *k* < *N seed* sequences (default: *k* = 50) consisting of the longest sequence, the most dissimilar sequence to it, and *k* – 2 randomly selected sequences.
3. Build a tree on seed sequences using UPGMA.
4. If there are non-seed sequences remaining:
  a. Assign non-seed sequences to their closest seeds to produce subclusters.
  b. Recursively apply the clustering procedure in each subcluster.
  c. Combine the resulting child trees with the current tree.
5. Return the tree.

While the default PartTree uses a fast, 6-mer-based dissimilarity measure, there are also modes performing more accurate alignment-based sequence comparisons: FastaPartTree and DPPartTree.

The random selection of seed sequences in PartTree results in the large instability of resulting trees—the topologies are sensitive to any variation, including reordering of the input sequences. The FAMSA2 medoid trees algorithm reduces stochasticity at the seed selection stage by performing a Partition Around Medoids (PAM) clustering and using the resulting cluster centers (medoids) as seed sequences. The remaining clustering steps are the same as in PartTree, with the exception that single linkage subtrees are used by default instead of UPGMA (UPGMA and NJ subtrees are also supported).

PAM^19^ is an optimization procedure that selects *k* medoids (representatives) from *N* points by minimizing the cost defined as the sum of dissimilarities between non-medoids and their closest medoids. As the algorithm takes all pairwise distances as input, the preparation stage requires *O*(*N*^2^) time (we assume a dissimilarity can be computed in constant time). The canonical PAM algorithm is deterministic and consists of two steps:

1. BUILD: Selects initial medoids in *O*(*N*^2^*k*) time.
2. SWAP: Iteratively selects the best possible medoid/non-medoid swaps until no improvement is possible or the maximum number of iterations is reached, resulting in the total time of *O*(*ik*(*N-k)*^2^), with *i* being the number of iterations.

These time complexities are prohibitive for large *N*, hence the several optimizations of PAM that improve execution times at the cost of introducing stochasticity from sampling:

- CLARA^19^: Runs PAM on a random subsample of size *M (*2*k < M* < *N*). This results in a time complexity of BUILD and SWAP steps together of *O*(*ikM*^2^).
- CLARANS^20^: Replaces BUILD with a random initial medoid selection (*O*(*k*) time) and restricts the SWAP step to a random subset of *k*(*M-k*) possible swaps (maxneighbor parameter). maxneighbor is often set relative to the number of possible swaps resulting in *O*(*kM*^2^) clustering time.
- FastCLARANS^17^: Eliminates *O*(*k*) redundant computations in SWAP, further reducing the time complexity of the clustering algorithm to *O*(*M*^2^).

The medoid tree algorithm recursively divides sequences into *k* clusters until they contain no more than *M* sequences according to the following steps:

1. *Initialization.* Assign all *N* sequences to the initial cluster.
2. *Recursive clustering.* If cluster size > *M* (default: *M* = 2000):
  a. *Seed selection.* Determine *k* seeds (default: *k* = 100) using medoid clustering:
    i. Draw a random sample of *M* sequences from the cluster (including the longest sequence).
    ii. Perform FastCLARANS clustering (maxneighbor = 0.1×*k*(*M-k*)). The longest sequence is fixed as one of the medoids.
    iii. Return the resulting medoids as seeds.
  b. *Internal tree construction.* Construct a single linkage tree *T* on the seed sequences using Prim’s algorithm.
  c. *Assignment.* Assign each non-seed sequence to its closest seed, forming *k* subclusters.
  d. *Recursion.* Apply clustering procedure to each subcluster (go to step 2) to obtain child trees *T*_1_, …, *T*_*k*_.
  e. *Tree extension*: Extend *T* with the child trees *T*_1_, …, *T*_*k*_. Otherwise (*cluster size ≤ M):*
  f. *Bottom tree construction.* Compute a single linkage tree *T* on all sequences in the cluster using Prim’s algorithm.
3. *Termination*. Return *T* to the higher recursion level (or as the final guide tree on the top level).

Running the medoid tree on ≤ *M* sequences is equivalent to FAMSA2’s standard mode. To maximize the computational performance of the algorithm, the recursive clustering of subclusters at the top recursion level is parallelized. Deeper recursion levels are processed sequentially to limit thread proliferation during the divide-and-conquer process.

#### Complexity analysis

To estimate the number of recursion levels, we make the realistic assumption that at every recursion level, the ratio between the maximum and the average cluster size is upper-bounded by a constant *r.* Thus, at the top level the largest cluster has the size *N*×*r/k*, on the second level: *N*×*(r/k)^2^*, and on the *L-*th level: *N*×*(r/k)^L^* (we consider the pessimistic case when the largest cluster at a given level was created from the largest cluster at the previous level). Given that the recursion stops when the cluster size drops to *M*, we get *M* =*N*×*(r/k)^L^.* The number of recursion levels is then *L*=log_*k/r*_*(N/M)*, which, under a reasonable assumption that *r* < *k, is O*(log *N*).

The processing of a bottom cluster with size *m* ≤ *M* consists of constructing a tree with Prim’s algorithm (step 2f) and requires *O*(*m*^2^) time. The total number of elements in all bottom clusters is *N*. The pessimistic case is when elements are distributed across clusters of maximum size

*M*. Given that there are *N*/*M* such clusters, the overall time complexity of processing all bottom clusters is *O*(*MN*).

The processing of an internal cluster with size *m* > *M* involves several steps. The time complexities of seed selection (step 2a; *O*(*M*^2^)) and internal tree construction (2b; *O*(*k*^2^)) do not depend on *m and* sum up to *O*(*M*^2^). Since the number of internal clusters at a given recursion level is upper-bounded by *N*/*M*, performing steps 2a and 2b across all clusters in a level requires O(*MN)* time. In contrast, the time complexities of assignment (2c; *O*(*km*)) and tree extension (2e; O(*m*)) depend on the cluster size. As the total number of elements at a given recursion level is at most *N*, the time complexity of steps 2c and 2e across all clusters in a level is *O*(*kN*). Given that there are *O*(log *N*) recursion levels, the processing of all internal clusters requires *O*(*M N* log*N*) time.

Overall, the time complexity of the medoid tree algorithm is *O*(*M N* log*N*).

#### Single linkage guide tree calculation via Prim’s algorithm

FAMSA2 utilizes Prim’s minimum spanning tree (MST) algorithm^21^ for a single-linkage guide tree calculation. Prim’s algorithm partitions sequences into two sets: *M* (sequences already included in an MST) and *R* (sequences not yet added to the MST). For each sequence in *R*, the smallest dissimilarity value to any of the sequences from *M* is tracked. At each step, the sequence *r* from *R* with the minimum dissimilarity is selected (parallelized across threads). The selected sequence *r* is moved from *R* to *M*, and the dissimilarity values between *r* and all remaining sequences in *R* are calculated (this step is also parallelized). This procedure repeats until *R* is empty. Prim’s algorithm is highly parallelizable and requires only linear space with the number of sequences (*N*), making it well-suited for large datasets.

#### Dissimilarity measure

For each pair of sequences *A* and *B*, the dissimilarity is calculated as: indel(*A,B*)^0.75^/LLCS(*A,B*), where

- LLCS(*A,B*) is the length of a longest common subsequence for *A* and *B*;
- indel(*A,B*) is the minimal number of single-symbol insertions or deletions necessary to transform *A* into *B*. It can be easily derived from LLCS: indel(*A,B*) = |*A*| + |*B*| -2 LLCS(*A,B*). This updated formula introduces a 0.75 exponent (compared to 1.0 in FAMSA), which was confirmed to improve the robustness to the sequence divergence.

LLCS is calculated using the bit-parallel approach that compares 64 residue pairs simultaneously. FAMSA2 supports CPU SIMD extensions including AVX, AVX2, AVX512 (novel in FAMSA2), and NEON (novel in FAMSA2). For example, with AVX512, a single thread can perform eight 64-symbol comparisons concurrently. Given that many threads can operate at the same time, on a 64-core AVX512-equipped CPU, up to 32,768 residue pairs can be compared at the same time.

#### Length-based dissimilarity prefiltering

In Prim’s algorithm, before computing the exact LLCS between a selected sequence *r* and each sequence *x* in *R*, FAMSA2 estimates a best-case LLCS value assuming the shorter sequence is a subsequence of the longer one. If this estimate still results in a higher dissimilarity than *x*’s current minimum to *M*, the exact LLCS calculation is skipped. This heuristic eliminates a few percent of pairwise comparisons, providing a moderate speedup (**Fig. S1**).

#### Parallel profile-profile alignment

In FAMSA, alignment-stage parallelism was limited to cases where multiple profile pairs could be aligned independently of each other. However, due to the single-linkage guide tree structure, higher tree levels often involve aligning single sequences or small profiles with much larger ones in a sequential, chain-like manner, limiting the parallelism.

FAMSA2 introduces a parallel implementation of both profile-profile and profile-sequence alignments. It takes advantage of atomic types, lock-free operations, and actively waiting thread barriers to avoid thread sleep latency. This enables efficient use of multiple threads even during chained alignments, significantly accelerating the process, especially when alignment time becomes the dominant factor in the total runtime.

#### Dynamic refinement of partial profiles

In FAMSA, refinement was applied only to final profiles with up to 1,000 sequences, as it proved ineffective for larger alignments. In FAMSA2, refinement is used during intermediate steps, specifically when two profiles, each with fewer than 1,000 sequences, are aligned to a profile with more than 1,000 sequences. This ensures refinement occurs while profiles remain small enough for effective improvement, enhancing large-scale alignment quality.

#### Output alignment compression

Because of FAMSA2’s high computational speed, writing alignments to disk can become the primary runtime bottleneck for large datasets. For example, FAMSA2 aligning 3 million sequences (average row length of ∼136,000 residues), produced ∼408 GB of uncompressed output in 284 seconds, requiring sustained write speeds exceeding 1.4 GB/s—well beyond typical HDD performance (∼250 MB/s). FAMSA2 integrates parallel Gzip compression into the output process: alignment data are partitioned across threads, each compressing its blocks independently, allowing compression and writing to proceed concurrently and maintaining overall efficiency.

#### Built-in column trimming

In phylogenetic analysis, full MSAs are often trimmed to remove poorly conserved columns (uninformative sites), typically using tools like trimAl^22^. FAMSA2 includes a built-in, simplified version of this step: users can set a minimum non-gap fraction per column for retention in the output, reducing alignment size and improving downstream analysis efficiency.

### Benchmarking alignment accuracy

Seven multiple sequence alignment (MSA) tools—Clustal Omega v1.2.4^12^, FAMSA v1.1^13^, FAMSA v2.4.1, Kalign v3.4.1^23^, MAFFT v7.526^24^, Muscle5 v5.3^10^, and the T-Coffee’s regressive method v13.46.0.919e8c6b^11^—were benchmarked across three categories of evaluation: structure-based, phylogeny-based, and function-based benchmarks (datasets described below). Among scalable MAFFT modes PartTree and DPPartTree, the latter exhibited better accuracy and running times in the preliminary experiments, thus was selected for the final evaluation. T-Coffee’s regressive pipeline was run with MAFFT sparsecore as a bottom aligner. All runtimes were benchmarked on a workstation equipped with an AMD Epyc 9554 CPU (64 cores clocked at 3.1 GHz) and 1152 GiB (approx. 1237 GB) RAM. Unless otherwise specified, all tools were run using 32 threads. Exact command-line parameters are provided in **Table S9**.

#### Structure-based benchmarks

The structure-based benchmarks included two complementary datasets. First, we constructed an extended version of the HomFam^12^ v37.0 (extHomFam) dataset, following a previous study^13^. The dataset was expanded by incorporating additional protein families from Homstrad^25^ (2 December 2023 release) and sequences from Pfam v37.0^26^ (UniProt variant), resulting in 390 protein families, each containing between 206 and 2,979,588 sequences (median: 46,855), with 3–41 Homstrad reference sequences (mean: 6) per family. For each family, MSAs were generated using the tested tools, and accuracy was evaluated by comparing the aligned Homstrad reference sequences to the original structural alignments. Accuracy was measured using the sum-of-pairs (SP) and total-column (TC) scores (range: 0–100) via T-Coffee’s ‘aln_compare’ option. The SP score reflects the proportion of correctly aligned residue pairs, while the TC score measures the fraction of correctly aligned columns shared with the reference. To assess accuracy with increasing numbers of homologs, we selected 121 extHomFam families containing more than 100,000 sequences. For each family, 20 subsets were generated by incrementally adding between 100 and 100,000 non-reference homologs to the reference sequences, producing alignments of increasing size. Accuracy stability was measured as the difference in SP and TC scores between the first (smallest) and last (largest) subsets. To evaluate the impact of non-homologous sequences, 20 composite subsets were created from the same 121 extHomFam families. Starting with a subset of 731 reference sequences (i.e., reference sequences combined from all 121 families), non-reference homologous sequences were added equally from each family, resulting in subset sizes ranging from 731 to 12,100,731 sequences. MSAs were computed for each subset, and accuracy was assessed by calculating SP and TC scores for each family, followed by averaging across all families.

A well-known limitation of the benchmark described above is its reliance on a limited number of reference structures per family, reflecting the uneven experimental coverage of the Protein Data Bank (PDB). To complement this, we constructed an additional structure-based benchmark leveraging the AlphaFold Database Clusters^9^, in which all sequences have predicted structures. In these datasets, structural alignments based on AlphaFold models have been shown to achieve accuracy comparable to alignments based on experimental structures. To avoid introducing bias from structure-based alignment methods, we assessed accuracy using the Local Distance Difference Test (LDDT)^16^, which estimates structural consistency based on the implied superposition of aligned residues. Accordingly, metadata for 24,963 clusters containing at least 1,000 proteins each were retrieved on March 6, 2025. PDB structures for 119,209,511 proteins were downloaded from the AlphaFold Protein Structure Database^8^. The clusters were divided into 13 size groups, ranging from 1,000 to 679,681 members. For accuracy assessment, 100 clusters were randomly sampled from each size group. For each sampled cluster, 100 sequences were randomly selected in 10 replicates. Columns containing only gaps were removed to obtain a refined 100-sequence MSA, which, along with the corresponding AlphaFold PDB structures, was used as input for FoldMason v7bd21ede^16^ to compute LDDT score, ranging from 0 to 1. The LDDT scores were averaged across the 10 replicates to provide a single accuracy measure for each cluster.

#### Phylogeny-based benchmarks

Phylogenetic accuracy was evaluated using two complementary benchmarks. They reflect the evolutionary assumptions underpinning MSA, where each column is interpreted as a statement about the shared ancestry of the residues it contains. While structural information is often used to assess alignment quality, it is an indirect proxy and may not fully capture evolutionary correctness^27^. In contrast, phylogenetic assessments aim to evaluate MSAs in terms of their capacity to preserve evolutionary signals. However, due to the lack of reference phylogenies for real data, such assessments rely on either simulations or indirect inference.

First, we used simulations to construct a dataset of 1,860 reference MSAs with known phylogenies. These were generated using the AliSim tool from IQ-TREE v2.4.0^28^ under diverse conditions, varying the number of sequences (1,000–100,000), substitution models (LG, JTT, WAG), sequence lengths (400–2,000 residues), sequence identities (8%–75%), and gap fractions (0%–99%). Unlike the extHomFam dataset, where alignment accuracy depends on a subset of reference sequences, the SP and TC scores were calculated based on all sequences within each reference MSA. In addition, we computed the expansion score, defined as the ratio of the number of columns in the tool-generated MSA to the number in the corresponding reference MSA, with values closer to 1 indicating better agreement with the reference MSA length (and typically, higher phylogenetical accuracy).

Second, we aimed to assess whether alignment quality affects the consistency of downstream phylogenetic inference. In practice, multiple sequence alignments are primarily used for phylogenetic reconstruction. While maximum likelihood (ML) methods are generally considered the most accurate, they are computationally intensive. Simpler distance-based methods, such as minimum evolution (ME), provide faster but typically less accurate estimates. However, these two approaches differ in their sensitivity to MSA errors: ML methods consider each alignment column individually, while distance-based methods can average over alignment inaccuracies. Higher-quality alignments should yield greater consistency between ML and ME trees^29^. To test this, we used 121 of the largest families from the extHomFam dataset. Eight families were excluded due to the presence of nonstandard amino acids, which caused failures in phylogenetic tree construction; consequently, 113 protein families were retained for phylogenetic analyses. For each family, we sampled 100,000 sequences and generated alignments using all tested tools. From each alignment, five replicate sub-alignments (MSA projections) of 100 randomly chosen sequences were extracted, preserving all alignment columns except those containing only gaps.

For each MSA projection, ML consensus trees were inferred using IQ-TREE with 1,000 ultrafast bootstrap replicates:

~~~
iqtree -s <alignment> -B 1000
~~~

ME consensus trees were reconstructed using FastME v2.1.6.4^30^ with BioNJ distance estimation, followed by 100 non-parametric bootstrap replicates:

~~~
fastme -i <alignment> -o <tree output> -m BioNJ -p LG -g 1.0 -s -n –
z 5 -b 100 -B <replicates file> -O <distance matrix output>
fastme -i <distance matrix> -s -n -z 5
~~~

Topological agreement between ML and ME consensus trees was measured using the Robinson–Foulds (RF) distance, which quantifies the number of differing bipartitions between two tree topologies. To obtain a normalized similarity score, the RF distance was divided by the maximum possible RF distance (max_RF) for the given trees. The topological agreement, expressed as a percentage, was then calculated as:

\text{Topological agreement} = \left(1 -\frac{\text{RF}}{\text{max_RF}}\right) \times 100

Agreement values were averaged across replicates and families for each subset size, yielding five topological agreement values per tool.

#### Function-based benchmarks

In addition to structural and evolutionary constraints, protein sequences encode functional information such as active sites, whose correct alignment is essential for downstream functional annotation. While structural and evolutionary benchmarks may indirectly reflect the alignment of conserved functional sites, convergent evolution and localized functional constraints may introduce confounding effects. To directly assess alignment accuracy in functional regions, we constructed a dedicated function-based benchmark focused on the conservation of active sites.

The benchmark was built from 772 enzyme domains annotated with 1376 active sites in Pfam v37.0. Protein sequences containing these domains were downloaded from UniProt, and active site residues were predicted using Pfam’s ‘pfam_scan.pl’ tool v1.6^26^ with the active site database ‘active_site.dat’ v37.0. For each protein, the sequence corresponding to the annotated domain was extracted from the full-length protein sequence. Domain sequences were excluded if the annotated domain was shorter than 25% of the length of the hidden Markov model domain or if more than 10% of residues were non-standard amino acids. The resulting domain families contain 1–11 active sites per family (median: 1), ranging in size from 11 to 1,515,446 sequences (median: 29,596), with 0.1% to 100% of sequences having known or predicted active sites (median: 86%).

Alignment accuracy within functional regions was evaluated by computing an active site entropy for each annotated site. For a given active site, the distribution of annotated residues across alignment columns was recorded. Let *p*_*i*_denote the fraction of active site residues aligned to column *i*. The entropy was defined as:

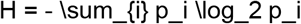

Lower values indicate higher concentration of active site residues within a single column (ideal alignment yields *H* = 0). Entropy scores were calculated for all 1,376 active sites and averaged to yield a single accuracy score for each MSA tool.

## Data Availability

Data is available at https://github.com/refresh-bio/FAMSA

## Code Availability

FAMSA2 is available at https://github.com/refresh-bio/FAMSA.

## Extended data figures

**Extended Data Figure 1.**
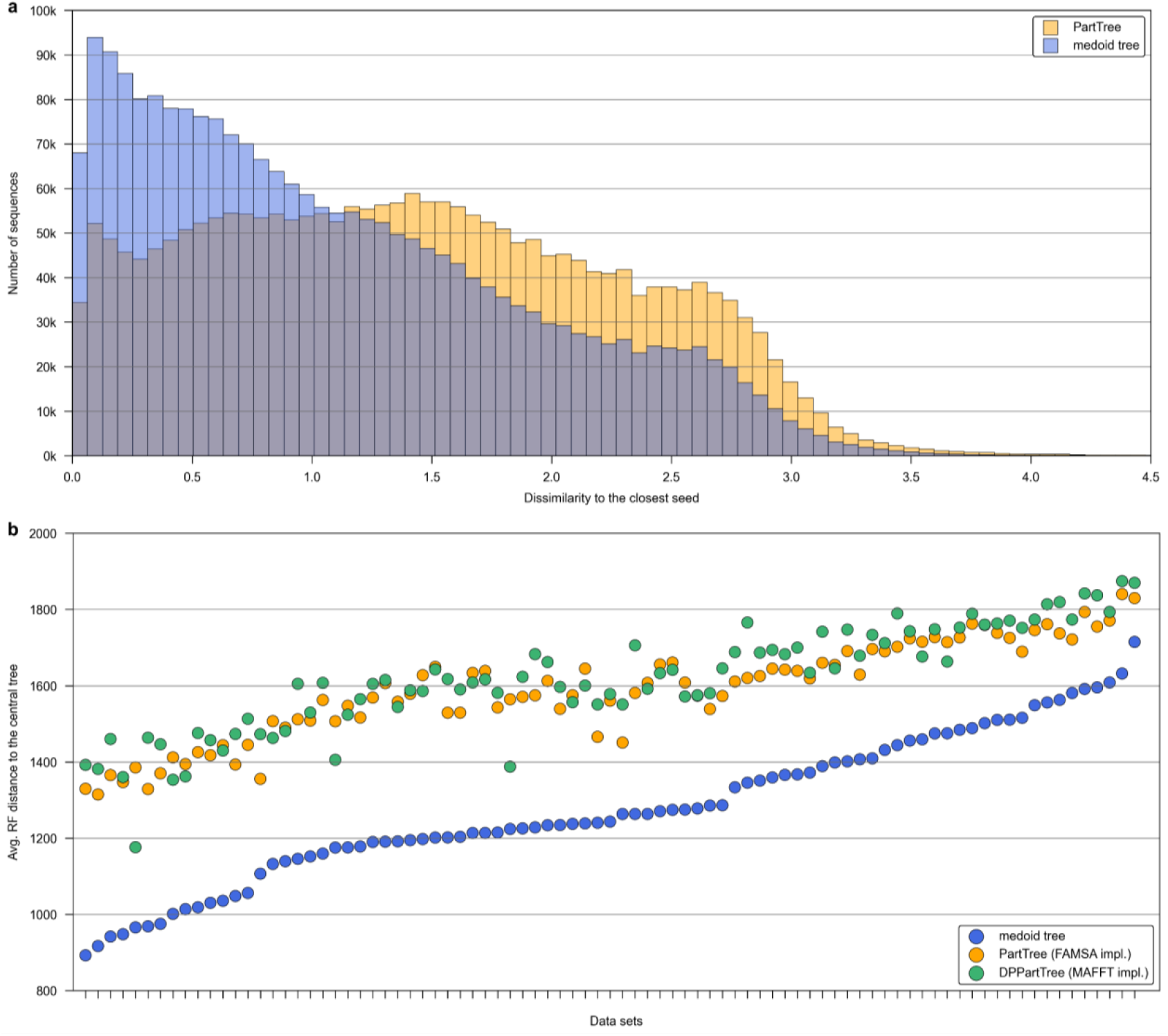
Properties of medoid trees and its alternative, PartTree from the MAFFT package, evaluated on extHomFam families with 15k to 40k sequences. **a,** Distributions of dissimilarities between sequences and their closest representatives (seeds) aggregated across all families within benchmark. The visualization concerns only the top recursion level with 50 seeds. Lower dissimilarities to representatives confirm medoid trees to provide better grouping than PartTree (reimplemented in the FAMSA framework to ensure consistency in dissimilarity measure). **b**, Stability of guide trees rendered by medoid tree, FAMSA-implemented PartTree, and MAFFT DPPartTree (PartTree with accurate dynamic programming-based dissimilarity measure). The stability of trees was measured per family (data set) with the following protocol: (i) randomly select 1,000 *core* sequences, (ii) generate 50 replicates, each with 5,000 sequences including all core sequences, (iii) generate a guide tree for every replicate and prune non-core sequences, (iv) calculate Robinson-Foulds (RF) distances between replicate trees, (v) determine central tree as the one minimizing the sum of RF distances to all other replicate trees.

**Extended Data Figure 2.**
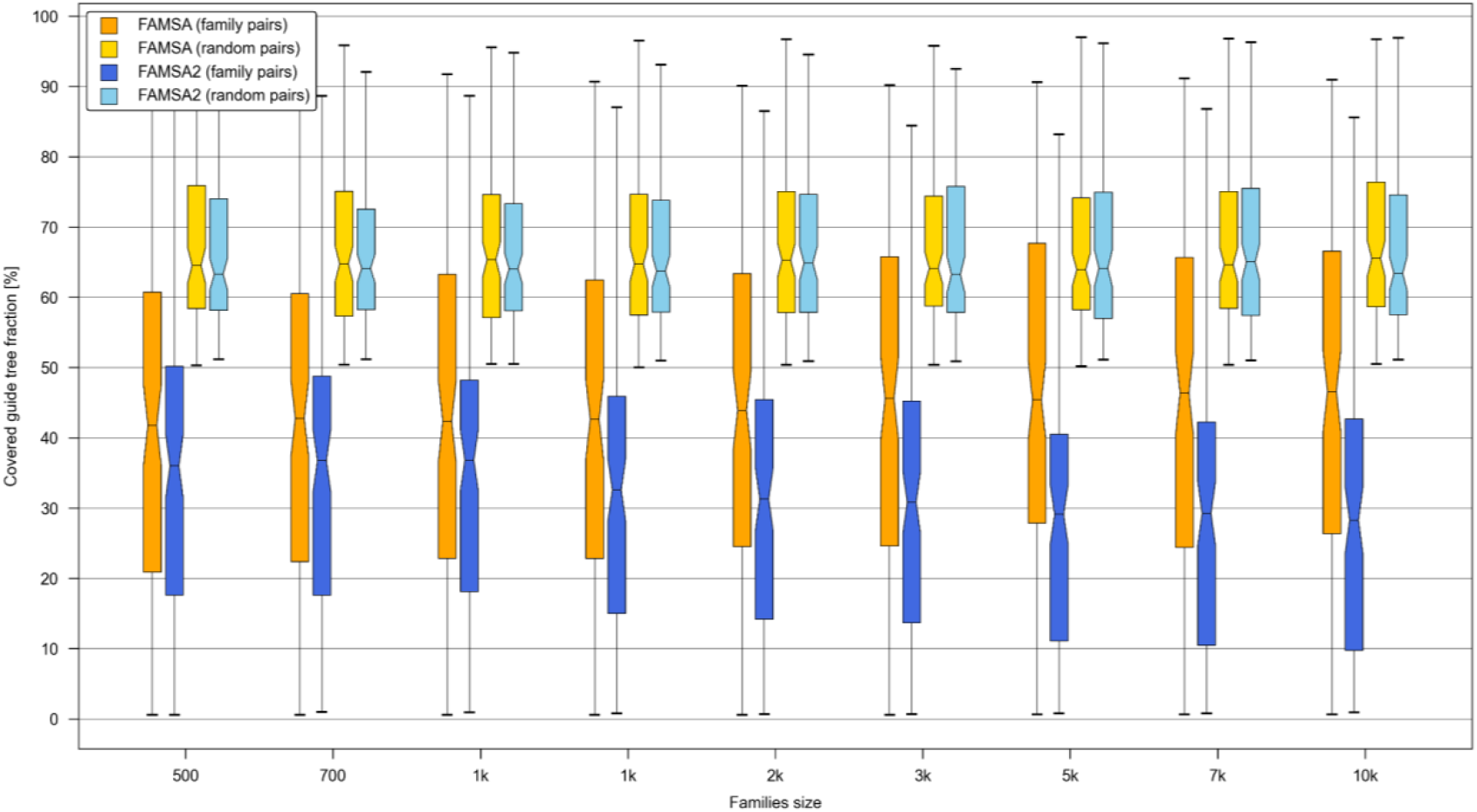
Comparison of LCS-based dissimilarity measures in FAMSA and FAMSA2 guide trees in terms of clustering homologous sequences. Guide trees were generated by FAMSA and FAMSA2 using concatenated sets of 121 extHomFam protein families with increasing total numbers of sequences. Each guide tree thus contained proteins from all 121 families. To assess the ability to cluster homologous sequences, 10,000 protein pairs were randomly sampled (with replacement) from each family, and the size of the smallest subtree containing each pair (defined by their last common ancestor) was determined. This size was expressed as a percentage of the total number of sequences in the guide tree, indicating how dispersed the pair was within the tree. Percentages were averaged over the 10,000 bootstrap samples per family, and the resulting 121 values (one per family) were visualized as boxplots. Across all dataset sizes, FAMSA2 dissimilarity measure produced significantly smaller subtrees than FAMSA (one-sided paired t-test, *p* < 0.001), indicating improved clustering of homologous sequences. As a negative control, 10,000 random protein pairs were sampled from the entire tree regardless of family, yielding no significant difference between methods.

**Extended Data Figure 3.**
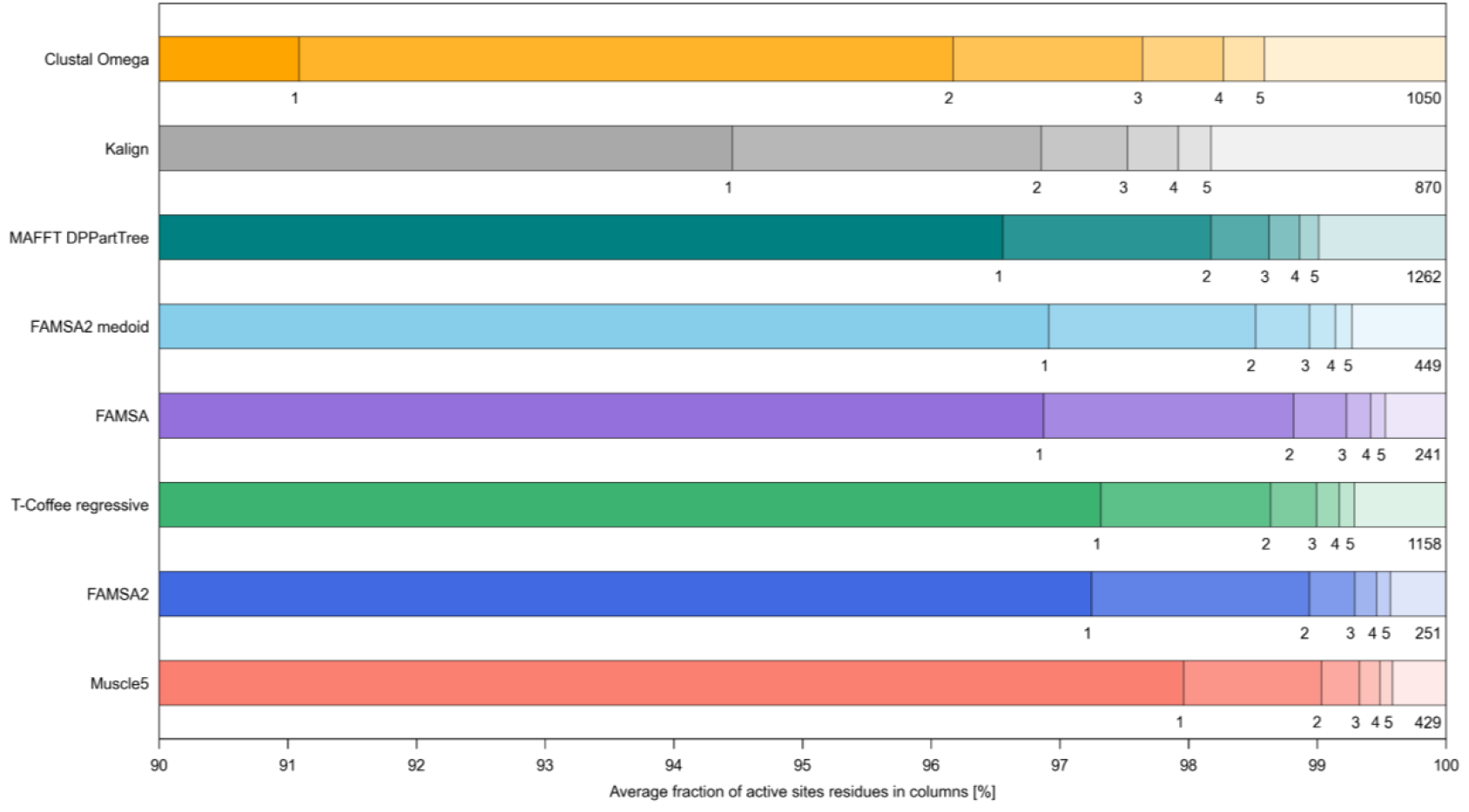
Concentration of active site residues in alignment columns across 772 Pfam enzyme families. For each active site, we established a distribution of residues across columns, numbered from most to least populated. These distributions were then averaged over all 1376 active sites for each tool and visualized as bars. For clarity, only the five most populated columns (labeled 1–5) are distinguished. The number on the right indicates the maximum number of distinct alignment columns over which an active site was distributed by a particular tool.

**Extended Data Figure 4.**
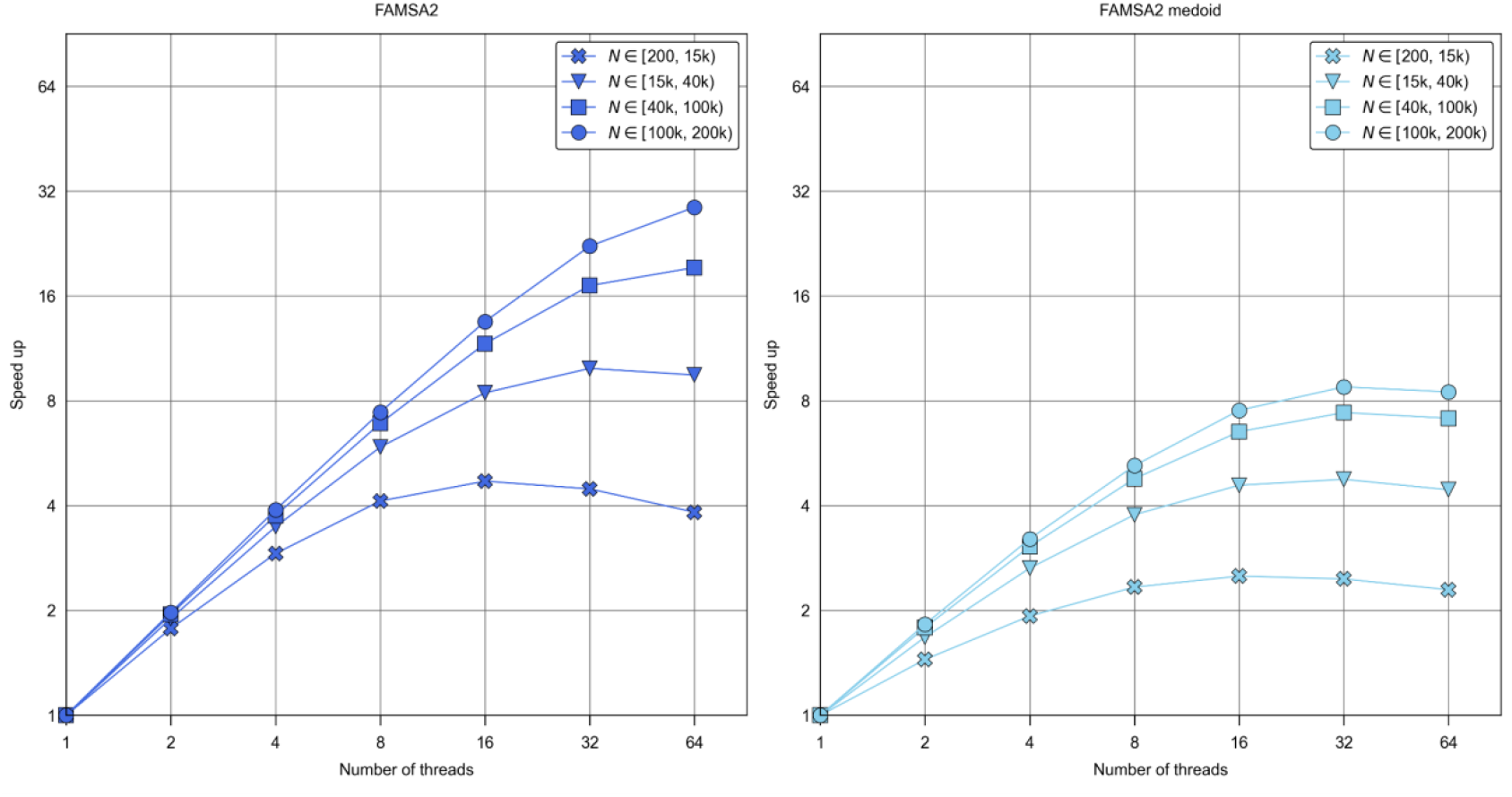
Scalability of FAMSA2 and its medoid guide tree variant across different numbers of threads on extHomFam protein families. Speedup (defined as the factor of acceleration relative to single-threaded execution) of **a,** FAMSA2 and **b,** FAMSA2 medoid tree as a function of the number of threads (1–64) on protein sequence sets from the extHomFam benchmark. Analyses were performed on extHomFam protein families grouped by size. FAMSA2 shows near-linear speedup up to 16 threads for larger datasets, achieving over 30-fold acceleration at 64 threads for the largest families, while the medoid tree variant exhibits more limited scalability, saturating around 8-fold speedup. These results demonstrate FAMSA2’s efficient multithreading on large-scale protein alignments.

## Extended Data Tables

**Extended Data Table 1. Summary of benchmarking results presented in Figure 2.** Each tool was evaluated across seven alignment scenarios: large protein families from extHomFam (SP/TC; Fig. 2a,b), increasing-size homologous datasets (SP/TC at largest size; Fig. 2c), mixed homologous and non-homologous datasets (SP/TC at largest size; Fig. 2d), structural consistency of AlphaFold clusters (LDDT; Fig. 2e), simulated MSAs (SP/TC and expansion; Fig. 2f), guide tree reconstruction (SP/tree agreement; Fig. 2g), and functional conservation of active sites (entropy; Fig. 2h). Accuracy metrics for Fig. 2c and 2d correspond to the final dataset size. A dash (–) indicates that the tool did not complete the corresponding test. The average rank column shows the mean rank for each method across metrics with numerical results from all methods.

## Supplementary Tables

**Table S1. Performance comparison of multiple sequence alignment tools on 390 protein families from the extHomFam v37.0 database.** Protein families are grouped by size and include the total and reference number of sequences. For each family and tool, runtime (seconds), peak memory usage (GB), and alignment accuracy (SP and TC scores, in %) are reported.

**Table S2. Average SP and TC scores for progressively expanded subsets of 121 protein families from the extHomFam v37.0 database.** Each family was stratified into 20 datasets by incrementally adding 0 to 100,000 homologous sequences to the reference alignment. Multiple sequence alignments were performed for each subset, and SP and TC scores were computed relative to the reference alignment and averaged across all families for each dataset size. The first column indicates the number of homologs added per family.

**Table S3. Evaluation of alignment robustness to protein family contamination with non-homologous sequences.** To simulate contamination, composite datasets were constructed by merging 731 reference sequences (one per each of the 121 largest extHomFam families) and progressively adding equal numbers of non-reference homologs from each family. This mixing introduces non-homologous sequences into individual family alignments, creating increasingly challenging conditions. Subset sizes range from 731 to over 12 million sequences. Multiple sequence alignments were generated for each subset using the listed methods. Alignment accuracy was evaluated by calculating SP scores separately for each of the 121 families—comparing aligned reference sequences within the composite MSA to their known reference alignment—and then averaging these scores across all families. “NT” (not tested) indicates that a method either exceeded a 36-hour runtime limit or had previously crashed at a smaller dataset size. Execution-related notes are summarized in the final row.

**Table S4. Structure-based alignment accuracy on 1166 AlphaFold protein clusters.** Each row corresponds to a single protein cluster, with columns summarizing cluster characteristics and the LDDT scores obtained using MSAs generated by alignment tools. For each cluster, 100 sequences were randomly selected in 10 replicates. Gap-only columns were removed from the resulting MSAs, which were then used with the corresponding AlphaFold structures as input to FoldMason to calculate LDDT scores (0–1). Reported values are the average LDDT scores across the 10 replicates and reflect the structural alignment accuracy achieved by each tool.

**Table S5. Running times and peak memory usage of multiple alignment tools executed on a sample of 390 AlphaFold clusters stratified by cluster size.** For each cluster size range, the table reports the number of clusters, total execution time (in seconds), and peak memory usage (in GB) for each tool. This subset (30 clusters per size range) was selected for runtime evaluation since running all tools on the complete set of 1,166 clusters used for accuracy assessment (see Table S4) was impractical on a single machine. Runtime measurements were performed on a single workstation.

**Table S6. Alignment accuracy and expansion of multiple sequence alignment tools evaluated on 1,860 simulated reference MSAs.** Accuracy is reported as SP and TC scores [%], and alignment expansion is defined as the ratio of the number of columns in the tool-generated MSA to the reference MSA. Summary statistics include the number of sequences, gap content [%], sequence identity [%], and sequence length range.

**Table S7. Topological agreement between maximum likelihood (ML) and minimum evolution (ME) trees for projected sub-alignments of 113 large extHomFam protein families.** Tree topology agreement was used as an indicator of alignment quality (Methods). For each of the 113 families from the extHomFam dataset, five subsets of 100,000 sequences were aligned with each tool, and five random sub-alignments of 100 sequences were extracted per alignment. These sub-MSAs preserved all columns except those containing only gaps. Topological agreement was quantified using normalized Robinson–Foulds (RF) similarity between consensus ML and ME trees. Higher agreement indicates better alignment accuracy. Mean SP scores for the corresponding full alignments are shown for reference.

**Table S8. Active site entropy across 1,376 sites in 772 Pfam domain families.** The characteristics of investigated active sites (e.g., Pfam accession, number of sequences, percentage of sequences with active site) together with accuracy of investigated tools measured as the entropy of the distribution of active sites’ residues across alignment columns.

**Table S9. Commands and versions of multiple sequence alignment tools used in this study.** Placeholders in angle brackets indicate input/output file names: <input> and <output> refer to FASTA files, and <guide_tree> to the guide tree file used in T-Coffee regressive alignments.

## Notes

https://github.com/refresh-bio/FAMSA

### Competing Interest Statement

The authors have declared no competing interest.

