## Supplementary figures and images for "FAMSA2 enables accurate multiple sequence alignment at protein-universe scale"

### Supplementary Fig S1

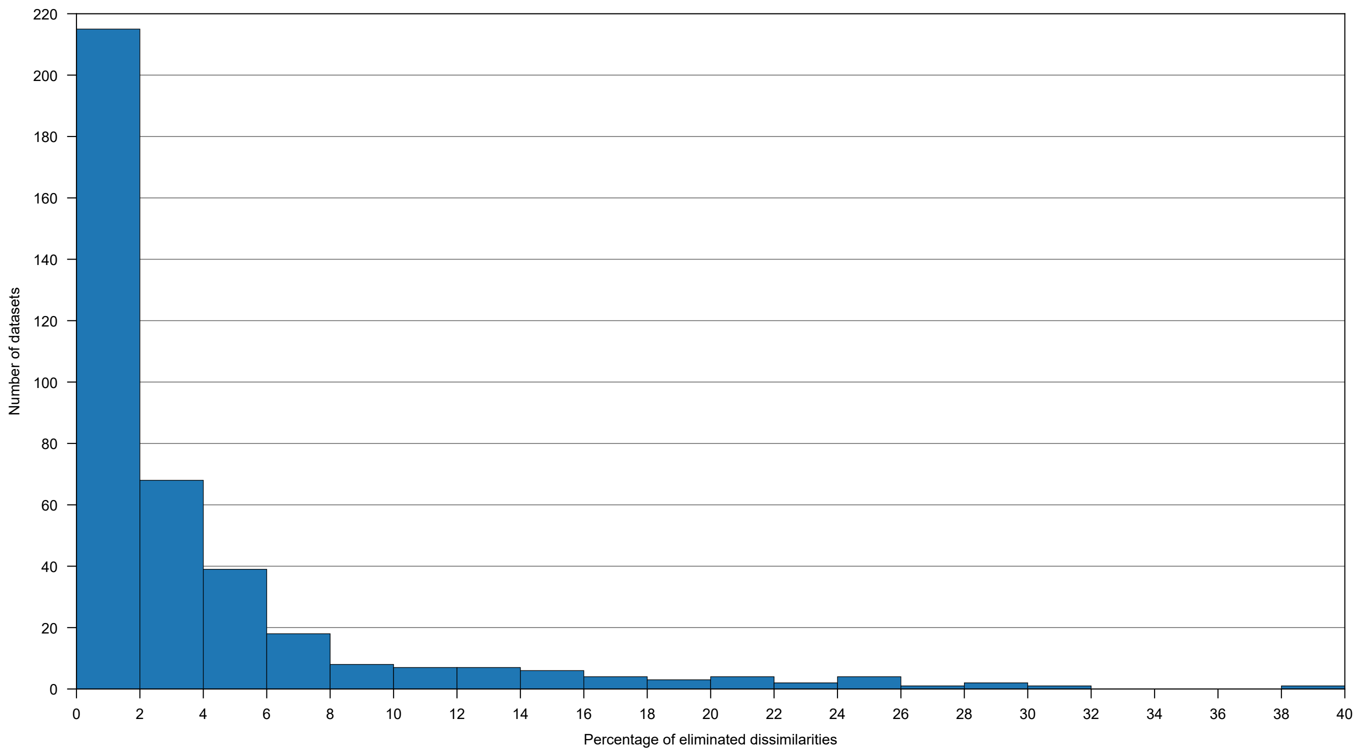
